# Reproductive fluids enabling cryptic female choice of paternity do not induce concomitant gamete-mediated paternal effects in embryos of hybridizing salmonid fishes

**DOI:** 10.1101/2024.07.09.602641

**Authors:** Tyler H. Lantiegne, Ranjan Wagle, Craig F. Purchase

**Affiliations:** Department of Biology, Memorial University of Newfoundland; St. John’s, Newfoundland & Labrador, A1C 5S7, Canada

## Abstract

Post-mating sexual selection in the form of cryptic female choice provides opportunities for females to bias paternity to favor preferred males. However, little is known regarding how cryptic female choice might affect offspring outside of paternity, via female modified changes to sperm environmental experience. Gamete-mediated paternal effects are widespread, and female alteration of sperm experience may play an unrecognized role in shaping cryptic female choice. Using hybridizing salmonid fishes that have documented female mediated conspecific sperm preference via differential upregulation of sperm swimming performance, we created artificial split-brood and split-ejaculate fertilizations to determine if sperm experience in different conditions influences offspring development. Prior to contact with eggs, sperm experienced 20s of swimming in either water, or water with the addition of conspecific ovarian fluid or heterospecific ovarian fluid. Over 186 days, we quantified hatch timing, hatchling size, and developmental stage and found that differential sperm experience created biologically irrelevant (average effect size of 1.06%) changes on offspring development, which were much smaller than the effects of hybridization itself (average effect size of 10.45% for the species of the father). Since ovarian fluid drastically changes sperm experience when compared to water, we conclude that females can use ovarian fluid to bias paternity without concomitant consequences to offspring development.

## Introduction

Polyandry occurs when females mate with several males in a given reproductive episode. This potentially allows eggs to be fertilized by higher quality fathers, which increases the chance of producing more fit offspring (Cothran, 2008; Garcia-Gonzalez & Simmons, 2005; Gowaty et al., 2010; Klemme et al., 2008). Polyandry also opens avenues of post-mating sexual selection. Among males, post-mating pre-zygotic sexual selection manifests as sperm competition where ejaculates from different individuals compete to fertilize the same eggs (Birkhead & Pizzari, 2002; Parker & Pizzari, 2010). In addition to a rich array of pre-mating behaviors, females can bias the outcome of this competition post-mating, through alterations of sperm behavior or experience, to favor certain males in a process referred to as cryptic female choice (Birkhead & Pizzari, 2002; Firman et al., 2017).

Cryptic female choice is taxonomically widespread, but mechanisms enabling it vary (Eberhard, 1996). Internally fertilized females can mechanically reject ejaculates (birds, Pizzari & Birkhead, 2000; Wagner et al., 2004) or deposit preferred sperm in more favorable locations within the reproductive tract (*Drosophila*, Manier et al., 2013). External fertilizers are limited to alterations of the aquatic sperm environment (Graziano et al., 2023) where female reproductive (Gasparini et al., 2020) fluids (egg water in aquatic invertebrates (Evans et al., 2012), ovarian fluid in fishes (Zadmajid et al., 2019)) change sperm swimming behaviour and influence sperm competition (Firman et al., 2017). Cryptic female choice changes paternity, which drastically alters the offspring’s genotype and therefore development. In contrast, paternal effects occur when offspring are influenced by the father independent of a change in gene sequence (paternity) and can occur due to adult alterations to sperm quality, or post-ejaculatory sperm experience (Evans et al., 2019; Purchase et al., 2021) - and can happen through changes in gene expression.

The examination of paternal effects is typically done through the lens of adult experience, e.g., diet (Evans et al., 2019; Purchase et al., 2021). However, there is increasing evidence that post-ejaculatory sperm environmental experience influences the development of offspring, which could also be considered paternal effects (Evans et al., 2019). For example, in internal fertilizers such as mice (Bromfield et al., 2014) and pigs (Robertson, 2007), variations in the environment of the female reproductive tract and subsequent chemical interactions between sperm and the environment, have effects on offspring survival and phenotype. In species with sperm storage such as kittiwakes, older sperm (sperm from chronologically older matings) have negative effects on hatch success and offspring survival and condition (Wagner et al., 2004; White et al., 2008). For external fertilizers, abiotic factors such as the temperature and pH of the fertilization environment (Byrne & Przeslawski, 2013; Kekäläinen et al., 2018; Lymbery et al., 2020), and the time sperm swim before fertilization (Alavioon et al., 2017, 2019; Crean et al., 2012) may influence offspring development. Since paternal effects can be altered by sperm environmental experience, and cryptic female choice drastically changes the fertilization environment to bias paternity, it follows that some mechanisms of cryptic female choice could affect individual sperm and lead to paternal effects. If cryptic female choice does impart concomitant paternal effects, these could be maladaptive and thus may constrain the mechanism or strength of cryptic female choice.

Given cryptic female choice changes paternity, which is generally compared among males of the same species, a conceptual default would be to test for paternal effects caused by cryptic female choice by examining female-male interactions within the same species, as has recently been done with a broadcast spawning marine invertebrate (Lymbery et al., 2020). However, a conceptually more powerful approach is to examine hybridizing species because the strength of modifications to sperm performance should be stronger across-species than across-individuals within-species. Polyandrous matings sometimes include males from different species (Holman & Kokko, 2013), and cryptic female choice allows females to bias paternity to favor males of their own species, through a process known as conspecific sperm preference (Howard, 1999). If cryptic female choice alters sperm in a way that induces paternal effects on offspring phenotype, we might expect to see relatively large effect sizes in hybridization situations because sperm are exposed to cues from a different species. In such context, differential development of hybridized eggs would result from some unknown combination of the genome of the father species, and paternal effects induced by sperm exposure to heterospecific female cues.

Conspecific sperm preference is seen in a wide range of taxa, including internal fertilizing terrestrial insects (Manier et al., 2013; Tyler et al., 2013) and birds (Cramer et al., 2016), and externally fertilizing fishes and invertebrates. It is comparatively easy for researchers to manipulate the fertilization micro-environment in external fertilizers, as parental effects can be more easily controlled. We therefore used sister species of external fertilizing fish that readily hybridize to test the hypothesis that cryptic female choice alters the development of offspring outside of the effects of paternity. To our knowledge, our study is the first examination of female induced gamete-mediated paternal effects in a vertebrate, and the first under hybridization situations in any taxa.

### Study system

Hybridization in salmonid fishes is common (Buss & Wright, 1958; Chevassus, 1979; Taylor, 2004). This is thought to be driven by high rates of polyandry (Weir et al., 2010) and alternative reproductive tactics in the form of sneaker males circumventing pre-mating female choice and male competition (Garcia-Vazquez et al., 2002; McGowan & Davidson, 1992; Weir et al., 2016). Some of the best studied are Atlantic salmon (*Salmo salar*) and brown trout (*S. trutta*). Hybrid matings and offspring from these species are common in the wild (Álvarez & Garcia-Vazquez, 2011), and hybrids can be easily produced with artificial fertilizations (Purchase et al., 2024) making them an ideal study species to examine the effects of cryptic female choice on offspring development.

In the genus *Salmo*, courtship behaviour helps females choose a preferred mate (Auld et al., 2019), but multiple unchosen males release sperm into the nest as she spawns (Esteve, 2005). In some of these situations, the unchosen males are of the heterospecific sister species. Large amounts of ovarian fluid are released with eggs and this dramatically alters sperm swimming behaviour. This has been reported to change swimming performance of conspecific sperm differently than heterospecific sperm, leading to conspecific sperm preference and most eggs being fertilized by males of her own species (Yeates et al., 2013). This cryptic female choice arises from a strong female modification of sperm experience and is a potential source of semen alteration that might influence the development of offspring (irrespective of paternity). Given the simple act of sperm swimming in water for different durations prior to fertilization has been reported to change the development of Atlantic salmon embryos (Immler et al., 2014), we predicted this mechanism of cryptic female choice (the species of ovarian fluid that sperm swim in) alters offspring development (and thus imparts gamete-mediated paternal effects) via changing the environment that sperm experience.

Fertilizing salmonid sperm always swim in the presence of some kind of ovarian fluid. Our primary goal was to test the hypothesis (1a) that when sperm are exposed to heterospecific ovarian fluid it induces different paternal effects than that of conspecific females, or alternatively (1b) salmon ovarian fluid consistently induces different paternal effects than trout ovarian fluid. Secondarily we tested (2) whether the evolution of ovarian fluid induces different paternal effects than occur in a water only swimming environment. To put results in context, we also hypothesized (3) that any influences of ovarian fluid type on paternal effects are similar for salmon and trout sperm, but modifications to offspring development resulting from sperm experience are less in magnitude (4) than those created by hybrid fertilization (changes in paternity of the sperm species).

## Methods

### Experimental design

The experiment tested the influences of ovarian fluid on offspring development from individual males. Any response was thus based on intra-ejaculate changes (a split-ejaculate design, Purchase & Rooke, 2020). We used a randomized block design whereby each block (n=6) contained semen (kept in isolation from one another) from one salmon and one trout. Critically, to reduce the influence of individual variation among females (Smith et al., 2019), ovarian fluid in each block came from a pool of equal proportions of three to four salmon and a separate pool of equal proportions from three to four trout. To trace offspring development, each block used a pool of equal proportions of eggs from three to four salmon, which were exposed to the sperm from the different treatments. Each traced group of sperm treated embryos were thus half-siblings (same father, one of three to four salmon mothers). The overall design thus combined split-ejaculates to control treatment, split-broods from pooled eggs to control individual variation in female quality, and a randomized block to allow for the collection of gametes and performing of artificial fertilizations on different days.

There were six treatments within each block (Table 1). These allowed us to determine if the medium in which sperm swim (sperm environmental experience) induces paternal effects and thus influences subsequent offspring development. Treatments SOF and TOF compared paternal effects on offspring from sperm that were exposed to salmon ovarian fluid (SOF, which was conspecific for salmon sperm (CSOF) and heterospecific (HSOF) for trout sperm) and trout ovarian fluid (TOF, heterospecific for salmon sperm and conspecific for trout sperm) for 20s prior to fertilization.

**Table 1:** Sperm experience treatments. Sperm from individual males were activated in 15 ml of swimming medium and then added to eggs after a 20s delay. Ovarian fluid was pooled among three to four females in each block to create species level effects. Legend: W=Water, SOF= Salmon ovarian fluid, TOF=Trout ovarian fluid, CSOF=Conspecific ovarian Fluid, HSOF=Heterospecific Ovarian Fluid, SS=Salmon sperm, TS=Trout sperm.

| **Treatment** | **Swimming Medium** | **Sperm Species** |
| --- | --- | --- |
| W-SS | Water | Salmon |
| SOF-SS | 33% Salmon Ovarian Fluid (CSOF) | Salmon |
| TOF-SS | 33% Trout Ovarian Fluid (HSOF) | Salmon |
| W-TS | Water | Trout |
| SOF-TS | 33% Salmon Ovarian Fluid (HSOF) | Trout |
| TOF-TS | 33% Trout Ovarian Fluid (CSOF) | Trout |

### Fish collection

Wild Atlantic salmon gametes were manually stripped from Exploits River fish in central Newfoundland, Canada (48.93 N, 55.67 W). Adults migrating upstream to spawn were trapped in a fishway on Grand Falls on September 7 2018, and transferred to indoor tanks (198 cm diameter, 91 cm depth) on September 30, where they were maintained under ambient conditions, similar to Rooke et al. (2020). Two weeks before gamete collection, all fish were measured for standard length and externally tagged to avoid repeat sampling. Gametes were collected over a period of 14 days (but all gametes were collected on the same day for a given block) beginning in early November. Salmon were anesthetized with MS-222, paper toweled dry, and stripped for semen into plastic bags and eggs into glass jars. Care was taken to prevent urine and feces contamination. Semen and eggs were insulated and stored on ice and were immediately transported to laboratory facilities at Memorial University for experimentation, which took place about 12 hours later.

Through coordinated efforts, for a given block, gametes were stripped from wild brown trout at “exactly” the same time as those from salmon, using similar methods. Spawning trout were captured by dipnet in tributaries of Windsor Lake near St. John’s Newfoundland, Canada (47.60 N, 52.78 W). These fish were introduced from Scotland in the 1880s (Westley & Fleming, 2011). Fish were anesthetized with clove oil, measured for length, stripped for eggs and sperm, and marked with a caudal fin clip to avoid repeat sampling of individuals. Although different anesthetics were used for both species, pre-gamete collection exposure to clove oil or MS-222 does not significantly affect sperm and egg function (Holcomb et al., 2004).

### Fertilization

Gametes were kept cool through all procedures, using a refrigerator and ice bath. To create the desired treatments (Table 1) pooled salmon and trout eggs were strained with an aquarium net to remove ovarian fluid (Purchase & Rooke, 2020), which was mixed with 5°C fresh water. Trout eggs were discarded after ovarian fluid solutions were made. Salmon eggs were then rinsed with 9psu salt water to remove any lingering ovarian fluid (Beirão et al., 2018). 300 µl of semen was used in each treatment. Based on previous experiences by our lab, the semen to egg ratio was low enough to avoid a ceiling effect (Beirão et al., 2018) but high enough to achieve a fertilization success that would produce adequate numbers of embryos for monitoring offspring performance.

The fertilization procedure followed Immler et al. (2014). Semen was put into the corner of a 50ml beaker and 5°C water was added, this mixture was then left to sit for 20s before being poured over ∼ 500 eggs. Eggs, semen, and the activating medium were gently mixed and left to rest for three minutes. Semen and activating medium were then removed from the eggs using an aquarium net as a sieve. Eggs were then covered in fresh water in static glass beakers and left overnight to water harden in a 5°C incubator.

Eggs that turned white during water hardening were deemed unviable and discarded the following morning. For each treatment (Table 1), 350 viable eggs were disinfected in a 1% ovadine solution, then separated equally into 7 PVC pipe incubation tubes (5.8 cm height x 5.8 cm diameter) with a screen on the bottom. Incubation tubes were transferred to one of three Marisource 4-tray vertical flow-through incubators. Each experimental block (n=6) was spread (Table 2) over 56 incubation tubes (6 treatments * 7 tubes * 50 embryos per tube) that were put into two incubation trays (each tray could hold 28 tubes). Salmon sperm and trout sperm tubes were split equally over both trays. In total there were 2100 viable eggs incubated per treatment (n=6), with the 12,600 individuals spread evenly over all blocks (n=6).

**Table 2:**
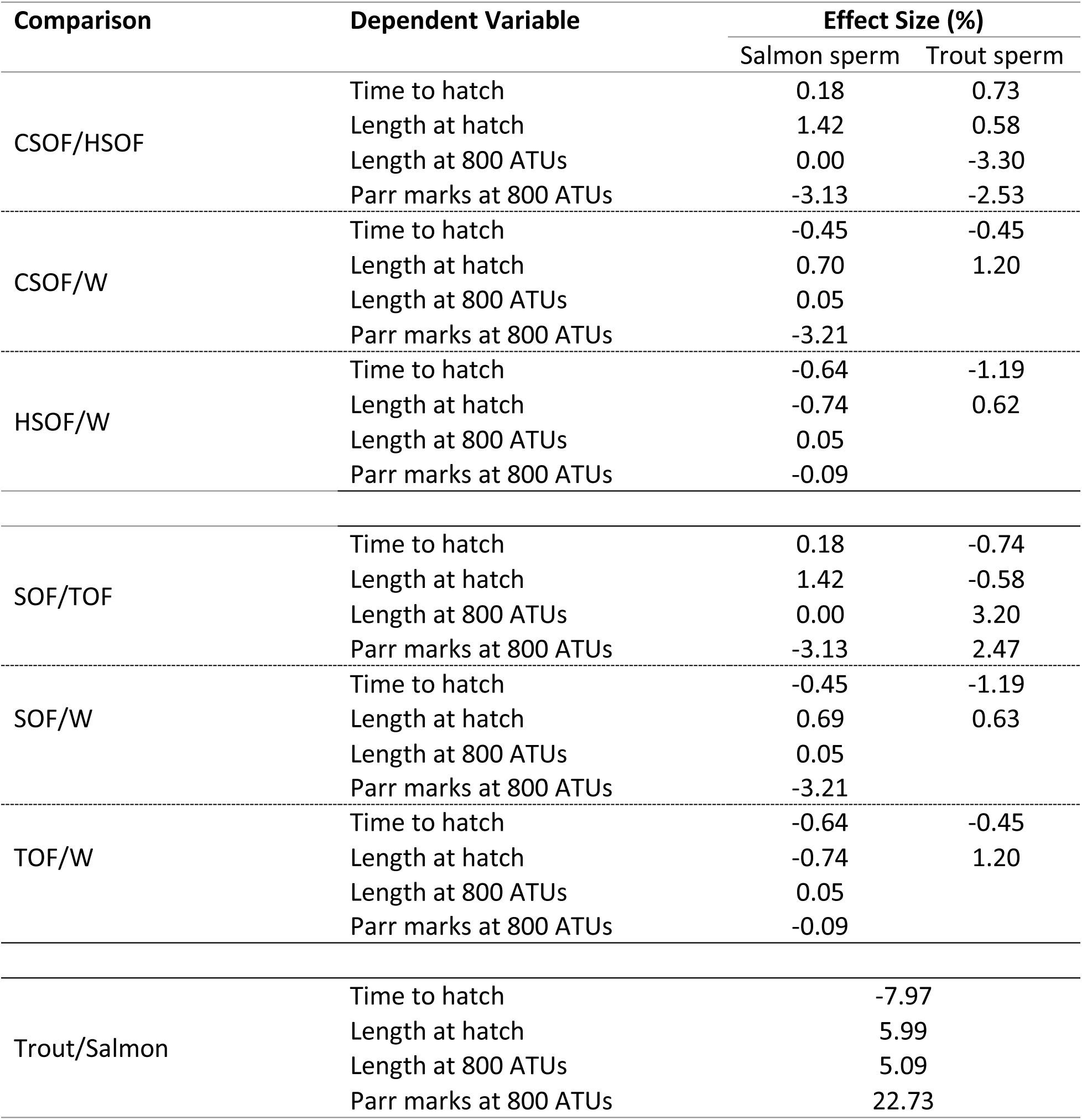
Effect sizes for sperm experience treatments and sperm species on the development of sibling embryos (Figure 1). Effect sizes were calculated using the means of the treatments (1-Latter/Former) or the sperm species values (1 - Salmon/Trout). Latter and former refer to order in the treatment column. Legend: W=Water, CSOF=Conspecific ovarian fluid, HSOF=Heterospecific ovarian fluid, SOF=Salmon ovarian fluid, TOF=Trout ovarian fluid. The average effect size for the species of the father (based on absolute values) was 10.45%, and the average effect size on the differential sperm experiences was 1.06%.

| Comparison | Dependent Variable | Effect Size (%) |  |
| --- | --- | --- | --- |
|  |  | Salmon sperm | Trout sperm |
| CSOF/HSOF | Time to hatch | 0.18 | 0.73 |
|  | Length at hatch | 1.42 | 0.58 |
|  | Length at 800 ATUs | 0.00 | -3.30 |
|  | Parr marks at 800 ATUs | -3.13 | -2.53 |
| CSOF/W | Time to hatch | -0.45 | -0.45 |
|  | Length at hatch | 0.70 | 1.20 |
|  | Length at 800 ATUs | 0.05 |  |
|  | Parr marks at 800 ATUs | -3.21 |  |
| HSOF/W | Time to hatch | -0.64 | -1.19 |
|  | Length at hatch | -0.74 | 0.62 |
|  | Length at 800 ATUs | 0.05 |  |
|  | Parr marks at 800 ATUs | -0.09 |  |
| SOF/TOF | Time to hatch | 0.18 | -0.74 |
|  | Length at hatch | 1.42 | -0.58 |
|  | Length at 800 ATUs | 0.00 | 3.20 |
|  | Parr marks at 800 ATUs | -3.13 | 2.47 |
| SOF/W | Time to hatch | -0.45 | -1.19 |
|  | Length at hatch | 0.69 | 0.63 |
|  | Length at 800 ATUs | 0.05 |  |
|  | Parr marks at 800 ATUs | -3.21 |  |
| TOF/W | Time to hatch | -0.64 | -0.45 |
|  | Length at hatch | -0.74 | 1.20 |
|  | Length at 800 ATUs | 0.05 |  |
|  | Parr marks at 800 ATUs | -0.09 |  |
| Trout/Salmon | Time to hatch | -7.97 |  |
|  | Length at hatch | 5.99 |  |
|  | Length at 800 ATUs | 5.09 |  |
|  | Parr marks at 800 ATUs | 22.73 |  |

### Incubation

Incubators were placed in individual sump tanks containing chillers set to 5°C, inside a temperature-controlled room at 9°C. The two blocks in each incubator experienced slightly different water temperatures due to variation between the three incubators, but importantly, treatments within a block were always exposed to the same temperature. Water in the tanks was partially changed weekly before hatch and every two days thereafter to maintain water quality. Temperature measurements of the tanks and the room were taken hourly, and water quality measurements (pH, nitrate, nitrite, and ammonia) were taken weekly. Recirculating water was passed through primary filtration, charcoal, and a UV sterilizer before entering the incubator at a flow rate of 16 L/min and gravity fed through the four incubation trays. Once the embryos reached approximately 20 accumulated temperature units (ATUs), any eggs/embryos that had turned white (Gaudemar & Beall, 1998) were removed weekly, except from 85-150 ATUs (∼ two to four weeks after fertilization), to minimize disturbances to living embryos during this vulnerable life-history stage when gastrulation occurs (Battle, 1944; Tang et al., 1987).

The goal was to compare gamete-mediated paternal effects induced by sperm exposure to different conditions. Given the size of the egg, maternal influences are generally more pronounced than those of the father early in development, the strength of these influences decreases with time (Eilertsen et al., 2008). Measurements thus represent a trade-off between experimental longevity and developmental stage. The experimental predictions are for offspring development, not fertilization success. To allow more time for paternal influence on offspring development (Eilertsen et al., 2008), a subset of eggs was checked at 240 ATUs (∼ six weeks after fertilization) for the presence of embryos (signifying cell division). This indicated that fertilization success was low in some treatments (semen volume was purposefully restricted and was the likely cause), but was not done in a comprehensive way to report fertilization success (any embryos taken early for fertilization success could not be used later to access development at a later stage). To compensate for low fertilization in some treatments and to prioritize standard developmental data collection across treatments before fish started to hatch at ∼ 350 ATUs (∼ ten weeks after fertilization), eggs were transferred from tube four and split equally into tubes one and seven (Table 2), to allow quantification of hatch times of embryos and an empty tube to store hatched individuals from other tubes. When embryos started to hatch within each treatment, individuals from tubes five and six from each treatment were counted daily and preserved in Stockard’s solution (Murray & Beacham, 1986). Embryos that hatched from tubes two and three were transferred into the empty tube four daily to track individual hatch dates and increase the number of individuals available for analysis at 800 ATUs. Data were not collected from hatchlings in tubes one and seven until they reached 800 ATUs. If hatch success was lower than 25% for a treatment within a block, then all embryos within that treatment were preserved at hatch. For the trout sperm W treatment, all embryos were taken at hatch due to low hatch success.

At hatch, salmonids are poorly developed and remain in gravel nests until ready to feed exogenously and emerge from the substrate (Mason, 1976). Unlike hatching, the timing of emergence can be somewhat subjective but generally occurs around 900 ATUs in Atlantic salmon (Gorodilov, 1996) as the endogenous yolk sac is almost completely consumed. We quantified development at 800 ATUs from fertilization (a precise number known for each embryo) which was ∼ nine to 10 weeks after first hatching and ∼ 22 weeks after fertilization. To reduce potential confounding variables on development post-hatch, hatchling densities were equalized at a maximum of 25 individuals per tube at approximately 575 ATUs (Supplement Table 1), roughly when all individuals completed hatching. Individuals were killed with an overdose of MS 222 at 800 ATUs and preserved in Stockard’s solution for later analysis. In total, the experiment ran for 186 days.

### Calculations and statistical analyses

The experiment was designed to test the influence of environmental conditions of sperm experience created by cryptic female choice, on the development of offspring. To assess development, we quantified eight metrics and report four in this manuscript: timing to 50% hatch, standard length at hatch, standard length at 800 ATUs, and the number of parr marks. These hatch metrics were chosen to replicate a previous study (Immler et al., 2014), while standard length at 800 ATUs was chosen to test for paternal effects at a longer timeframe than the previously cited study (which was three weeks). Parr mark count was chosen as an additional metric for development. An additional four metrics (head length, caudal ray count at hatch, proportion of embryos with separated adipose fins, caudal ray count at 800 ATUs), showed the same patterns as those reported, and are not included (see Lantiegne, 2021).

To calculate time to 50% hatch, each egg was given a binary number (0 = not hatched, 1= hatched) on each day. Hatch numbers were then used to create a logistic regression model relating day to hatch. Data collection stopped 1 week after the last embryo within a block hatched, as based on previous experiments in our lab, it was assumed that if no offspring hatched over a week, then no more offspring were likely to hatch. Equation 1 was solved for y = 0.5 to determine the day value where 50% of the offspring had hatched and then converted to ATUs (to ensure data was standardized across blocks).

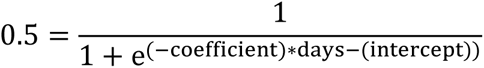

Equation 1: Logistic regression equation.

We used three additional metrics to quantify offspring development rate. Standard length at hatching, standard length at 800 ATUs from fertilization, and parr mark counts at 800 ATUs. Embryos were photographed with a Leica DFC420 camera mounted on a Leica M80 dissection microscope, and length measurements taken using ImageJ (Schneider et al., 2012). Effect sizes between treatments were created using mean values and the equation (1-X_2_/X_1_). Where X represents the means for the appropriate treatments being compared.

Several hypotheses were tested with the same modelling framework (Equation 2) whereby sperm species (salmon or trout), swimming medium (water only, conspecific ovarian fluid vs heterospecific ovarian fluid – or salmon ovarian fluid vs trout ovarian fluid) and their interaction were fixed effects, while block was a random effect. If the interaction was significant, the model was run separately for salmon and trout sperm. To test the primary hypothesis that cryptic female choice affects the development of offspring, we compared the development of siblings that were fathered by sperm exposed to conspecific ovarian fluid (SOF for salmon sperm, TOF for trout sperm) to those exposed to heterospecific ovarian fluid (TOF for salmon sperm, SOF for trout sperm). Alternatively, there maybe consistent trends created by each species of ovarian fluid, regardless of its relation to the father species. To test this, we repeated the same analysis as above, but recoded the ovarian fluid from being CSOF-HSOF (to the sperm) to TOF-SOF. Next, we tested the hypothesis that sperm experience in the absence of ovarian fluid (un-natural) affects development differently than experience in conspecific ovarian fluid.

Since Atlantic salmon and brown trout are very closely related, we hypothesized that the response of sperm experience would be similar for offspring sired by either species’ sperm which was evaluated using the interaction term in the model. Finally, we tested whether hybridization itself influenced development using sperm species term in the model.

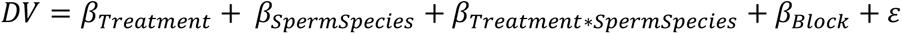

Equation 2: Model Equation. Data tested for each hypothesis was different depending on which treatments were compared. Error structure is not indicated because it varied for each model (see text).

We used a generalized linear mixed effects modelling approach to test for the effect of our treatments on offspring development. Models were tested using the ‘lmertest package’ (Kuznetsova et al., 2017) in R. Standard length at hatch and at 800 ATUs and time to hatch models were assumed to have a normal error structure and were tested using Type 3 ANOVA with Satterthwaite’s method. Parr mark count at 800 ATUs had a Poisson distribution, so tested using Wald Chi-Square tests. Due to poor hatch success, which resulted in no data on hatch length and parr mark count at 800 ATUs in the water treatment (W) of trout sperm (Father-trout), the fixed-effect model matrix was rank deficient, leading to the removal of the corresponding column/coefficient during model estimation (see Table 2 and Figure 1). Analysis of assumptions of parametric statistics revealed that our models for standard length at hatch and at 800 ATUs did not have a normal error structure. After transformation and other error structures did not meet this assumption, p-values around our α=0.05 at a threshold of p=0.025<x<0.075 were randomized to generate assumption free p-values (Ludbrook, 1994). All interactions, unless otherwise noted, were not significant at a threshold of α=0.05.

**Figure 1:**
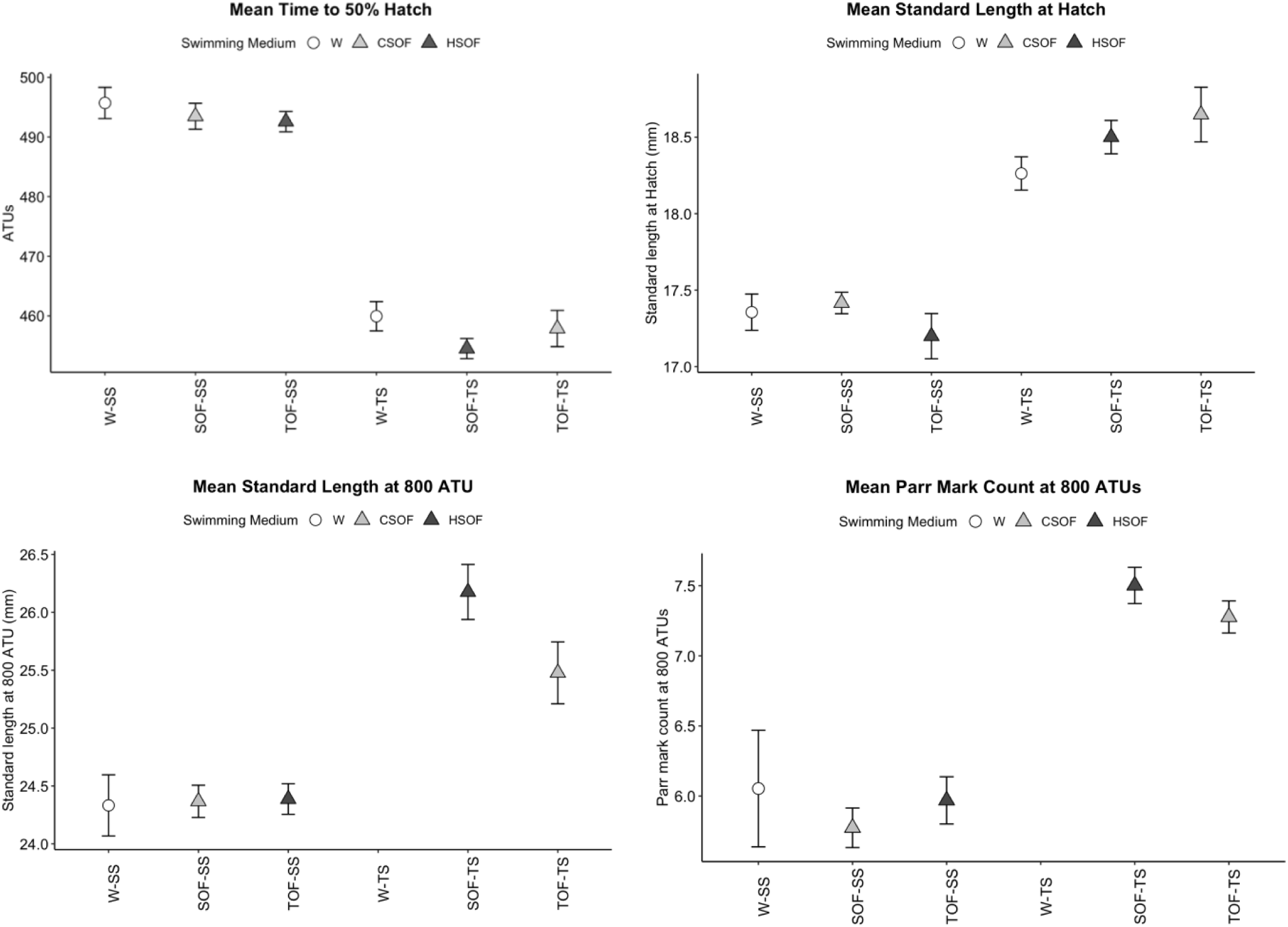
Developmental characteristics of Atlantic salmon (closed symbols) and hybrid salmon female (f) X brown trout male (m) embryos (open symbols) when sperm experienced different environmental treatments (Table 1) prior to contact with Atlantic salmon eggs. Each datum is the mean ± standard error among 6* blocks of parent fish. Panels: (a) accumulated temperature units (ATUs) to 50% hatch, (b) standard length at hatch, (c) standard length at 800 ATUs, (d) parr mark count at 800 ATUs. Legend: W=Water, SOF=Salmon ovarian fluid, TOF=Trout ovarian fluid. *Poor hatch success in some treatments resulted in incomplete data at 800 ATUs (means for treatments W-SS and TOF-SS to be taken from five blocks, while no data were available for W-TS). Blocks are represented with numbers.

## Results

We expected offspring development would be altered through sperm swimming in conspecific (SOF-SS, TOF-TS) vs. heterospecific ovarian fluid (SOF-TS, TOF-SS) and any pattern would be the same for salmon and trout sperm (Figure 1). As expected, this pattern and those that followed were the same for both salmon and trout sperm, but offspring development was only slightly altered (Table 2) by differential ovarian fluid experience.

At hatch, there was no significant effect of conspecific ovarian fluid vs. heterospecific ovarian fluid on time to hatch (Figure 1a, Table 3 rows 3, 4). For hatch size (Figure 1b), the ovarian fluid type was significant. Hatch size was significantly smaller when sperm were exposed to HSOF (Table 3 rows 9, 10), but the effect size (Table 2) was very small (salmon sperm = 1.42%, trout sperm = 0.58%). At 800 ATUs from fertilization (months after hatching), there were no significant influence of CSOF vs HSOF on parr mark count (Figure 1d, Table 3 row 21). The interaction was significant for body size at 800 ATUs (Figure 1c, Table 3 row 15). For salmon sperm, there was no effect (Table 3 rows 15, 16). Offspring from trout sperm were significantly (Table 3 rows 15, 16) larger (−3.30%) when exposed to HSOF. The very small effect sizes for trout sperm were in opposite directions for length at hatch vs length at 800 ATUs and are likely spurious.

**Table 3:** Results of statistical analysis for sperm experience treatments and sperm species for each variable. Full model refers to models that included both salmon and trout sperm. If significant species interactions were found, the data were broken, and the model re-run for salmon sperm and trout sperm separately. Due to poor hatch success in some treatments full models were not possible for all combinations. Legend: W=Water, CSOF=Conspecific ovarian fluid, HSOF=Heterospecific ovarian fluid. * represents values significant at α=0.05.

| Row no. | Dependent Variable | Independent Variables | Full model |  |  | Salmon sperm model |  |  | Trout sperm model |  |  |
| --- | --- | --- | --- | --- | --- | --- | --- | --- | --- | --- | --- |
| | | | F/$\chi^2$/t | df | P | F/$\chi^2$/t | df | P | F/$\chi^2$/t | df | P |
| 1 | Time to Hatch | FS*SM | 0.49 | 2, 23.12 | 0.622 |  |  |  |  |  |  |
| 2 |  | Father Species (FS) | 1015.24 | 2, 23.24 | <0.001* |  |  |  |  |  |  |
| 3 |  | Swimming Medium (SM) | 4.63 | 1, 23.12 | 0.020* |  |  |  |  |  |  |
| 4 |  | Tukey CSOF-HSOF | 1.54 | 23.1 | 0.290 |  |  |  |  |  |  |
| 5 |  | CSOF-W | -1.55 | 23.1 | 0.284 |  |  |  |  |  |  |
| 6 |  | HSOF-W | -3.04 | 23 | 0.015* |  |  |  |  |  |  |
| 7 | Length at Hatch | FS*SM | 3.02 | 2, 1426.7 | 0.05 |  |  |  |  |  |  |
| 8 |  | Father Species | 454.27 | 1, 1418.3 | <0.001* |  |  |  |  |  |  |
| 9 |  | Swimming Medium | 9.9 | 2, 1426.4 | <0.001* |  |  |  |  |  |  |
| 10 |  | Tukey CSOF-HSOF | 4.34 | 1427 | <0.001* |  |  |  |  |  |  |
| 11 |  | CSOF-W | 2.04 | 1426 | 0.101 |  |  |  |  |  |  |
| 12 |  | HSOF-W | -0.76 | 1427 | 0.726 |  |  |  |  |  |  |
| 13 | Length at 800 ATU | FS*SM | 23.71 | 1, 1700.0 | <0.001* |  |  |  |  |  |  |
| 14 |  | Father Species | 532.03 | 1, 1689.8 | <0.001* |  |  |  |  |  |  |
| 15 |  | Swimming Medium | 22.55 | 2, 1698.8 | <0.001* | 3.8 | 2, 1193 | 0.023* | 25.76 | 1, 451.1 | <0.001* |
| 16 |  | Tukey CSOF-HSOF |  |  |  | -1.147 | 1197 | 0.485 | -4.987 | 451 | <0.001* |
| 17 |  | CSOF-W |  |  |  | -2.747 | 1187 | 0.017* |  |  |  |
| 18 |  | HSOF-W |  |  |  | -1.898 | 1199 | 0.139 |  |  |  |
| 19 | Parr Count at 800 ATU | FS*SM | 0.78 | 1 | 0.376 |  |  |  |  |  |  |
| 20 |  | Father Species | 133.39 | 1 | <0.001* |  |  |  |  |  |  |
| 21 |  | Swimming Medium | 1.33 | 2 | 0.513 |  |  |  |  |  |  |
| 22 |  | Tukey CSOF-HSOF |  |  |  |  |  |  |  |  |  |
| 23 |  | CSOF-W |  |  |  |  |  |  |  |  |  |
| 24 |  | HSOF-W |  |  |  |  |  |  |  |  |  |

Ignoring conspecific vs heterospecific identification of ovarian fluid to sperm, and focusing solely on female species, time to hatching was not different if sperm swam in salmon vs trout ovarian fluid (Figure 1a, Table 4 rows 3,4). For size at hatch, the interaction term was significant (Table 4 row 7), and the model was run for each father species. Swimming medium (SOF-TOF) did not influence embryo development from trout sperm (Table 4 row 9). Salmon sperm exposed to SOF hatched significantly larger than those in TOF (Figure 1b, Table 4 rows 9, 10, but the effect size was very small 1.42% (Table 2). At 800 ATUs from fertilization, parr mark count was not influenced by SOF-TOF (Figure 1d, Table 4 row 21). The interaction was significant for body size at 800 ATUs (Table 4, row 13). For salmon sperm, SOF-TOF had no effect (Figure 1c, Table 4 rows 15, 16). Embryos developing from trout sperm hatched larger in SOF than those from TOF (Figure 1c, Table 4 rows 15,16) and the effect size was 3.20% (Table2).

**Table 4:** Results of statistical analysis for sperm experience treatments and sperm species for each variable. Full model refers to models that included both salmon and trout sperm. If significant species interactions were found, the data were broken, and the model re-run for salmon sperm and trout sperm separately. Due to poor hatch success in some treatments full models were not possible for all combinations. Legend: W=Water, SOF=Salmon ovarian fluid, TOF=Trout ovarian fluid. *represents values significant at α=0.05.

| Row no. | Dependent Variable | Independent Variables | Full model |  |  | Salmon sperm model |  |  | Trout sperm model |  |  |
| --- | --- | --- | --- | --- | --- | --- | --- | --- | --- | --- | --- |
| | | | F/$\chi^2$/t | df | P | F/$\chi^2$/t | df | P | F/$\chi^2$/t | df | P |
| 1 | Time to Hatch | FS*SM | 1.3 | 2, 23.12 | 0.292 |  |  |  |  |  |  |
| 2 |  | Father Species (FS) | 1015.23 | 2, 23.24 | <0.001* |  |  |  |  |  |  |
| 3 |  | Swimming Medium (SM) | 3.97 | 1, 23.12 | 0.033* |  |  |  |  |  |  |
| 4 |  | SOF-TOF | -0.891 | 23.1 | 0.651 |  |  |  |  |  |  |
| 5 |  | Tukey SOF-W | -2.771 | 23.1 | 0.028* |  |  |  |  |  |  |
| 6 |  | TOF-W | -1.848 | 23 | 0.177 |  |  |  |  |  |  |
| 7 | Length at Hatch | FS*SM | 12.18 | 2, 1422.5 | <0.001* |  |  |  |  |  |  |
| 8 |  | Father Species | 454.27 | 1, 1418.3 | <0.001* |  |  |  |  |  |  |
| 9 |  | Swimming Medium | 1.17 | 2, 1426.1 | 0.31 | 11.78 | 2, | <0.001* | 2.45 | 2, 435.08 | 0.086 |
| 10 |  | SOF-TOF |  |  |  | 4.37 | 987 | <0.001* |  |  |  |
| 11 |  | Tukey SOF-W |  |  |  | -0.172 | 986 | 0.984 |  |  |  |
| 12 |  | TOF-W |  |  |  | -3.57 | 991 | <0.001* |  |  |  |
| 13 | Length at 800 ATU | FS*SM | 42.56 | 1, 1699.1 | <0.001* |  |  |  |  |  |  |
| 14 |  | Father Species | 532.03 | 1, 1689.8 | <0.001* |  |  |  |  |  |  |
| 15 |  | Swimming Medium | 13.21 | 2, 1699.1 | <0.001* | 3.8 | 2,1193 | 0.023* | 25.763 | 1, 451.09 | <0.001 * |
| 16 |  | SOF-TOF |  |  |  | -1.147 | 1197 | 0.4851 | 4.987 | 451 | <0.001 * |
| 17 |  | Tukey SOF-W |  |  |  | -2.747 | 1187 | 0.017* |  |  |  |
| 18 |  | TOF-W |  |  |  | -1.898 | 1199 | 0.140 |  |  |  |
| 19 | Parr Count at 800 ATU | FS*SM | 0.43 | 1 | 0.512 |  |  |  |  |  |  |
| 20 |  | Father Species | 125.29 | 1 | <0.001* |  |  |  |  |  |  |
| 21 |  | Swimming Medium | 1.68 | 2 | 0.43 |  |  |  |  |  |  |
| 22 |  | SOF-TOF |  |  |  |  |  |  |  |  |  |
| 23 |  | Tukey SOF-W |  |  |  |  |  |  |  |  |  |
| 24 |  | TOF-W |  |  |  |  |  |  |  |  |  |

If ovarian fluid type is not important, we hypothesized that the absence of ovarian fluid (W) would create strong effects when compared to sperm swimming in conspecific ovarian fluid (W/CSOF). This was not the case (Figure 1). There was no effect on time to hatch (Figure 1a, Table 3 rows 3, 5) or size at hatch (Figure 1b, Table 3 rows 9,11). At 800 ATUs, trout sperm comparisons could not be made because too few offspring were available for the W (water) treatment. For salmon sperm there was no effect on parr mark count (Figure 1d, Table 3 row 21), length was significant (Table 3 rows 15, 17) but the effect size was extremely small (Figure 1c, Table 2).

As expected, hybridization (father species) influenced the development of embryos derived from salmon eggs more than paternal effects (sperm experience) did. In all metrics, father species was significant (Table 3) with greater effect size than any of the sperm swimming treatments (Table 2). When split-brood salmon eggs were hybridized by brown trout sperm they hatched 7.97% sooner (Figure 1a, Table 3 row 2) and at 5.99% larger sizes (Figure 1b, Table 3 row 8) and continued to show faster development to 800 ATUs (Figure 1c, length 5.09%, Table 3 row 14; Figure 1d, parr mark count 22.73%, Table 3 row 20), than if they had a salmon father.

## Discussion

When hybridization occurs, if development of hybridized eggs is different (common) than sibling eggs sired by conspecific sperm, it seems obvious that these changes are caused directly by the genome of the father species. However, the presence of female reproductive fluids means that the heterospecific sperm must swim in, and therefore have pre-fertilization experiences in, a foreign environment. Recent work has reported that sperm experience can substantially alter offspring development, and therefore it is possible that the altered development of hybrid offspring is some combination of the direct effect of the siring sperm’s genome, and the indirect effect of the exposure of that sperm to female reproductive fluids that regulate conspecific sperm preference.

We sought to determine if cryptic female choice influences offspring development independent of paternity by evaluating the effects of chemically/physically mediated conspecific sperm preference in the context of hybridizing fish. To our knowledge, this is the first time parental effects have been analyzed in the context of hybridization and the first time gamete-mediated paternal effects have been examined in vertebrates in the context of cryptic female choice. We found that sperm experience in conspecific ovarian fluid vs. heterospecific ovarian fluid did not strongly alter embryo development. More surprisingly, neither did sperm experience in congeneric ovarian fluid vs. the un-natural condition of water only. Previous work on these fish populations (Lantiegne & Purchase, 2023; Purchase & Rooke, 2020) has shown that sperm experience is heavily modified by exposure to ovarian fluid vs. water, but here we report no substantial effect in terms of non-genetic alterations to offspring. This implies that cryptic female choice is free to modify paternity without concomitant consequences to embryo development caused by gamete-mediated paternal effects.

As expected, hybridization did strongly influence development, as salmon eggs fathered by trout sperm hatched faster and larger than pure-species siblings. This pronounced difference continued to 800 ATU after fertilization (months after hatching), as hybrids were more developed. Our data supports that work of others, who have reported that trout (*S. trutta*) develop faster than salmon (*S. salar*), and that hybrids are intermediate (McGowan & Davidson, 1992b). Faster development of trout could be partially caused by sperm experience prior to fertilization, if trout ovarian fluid induces changes differently from salmon ovarian fluid. If this is the case, the predicted rates of offspring development from different cross types would be ranked (female-male) trout-trout, trout-salmon, salmon-trout, salmon-salmon. Data are available to support this prediction (McGowan & Davidson, 1992a; Poulos, 2019). However, our experiment suggests that faster development of trout-salmon over salmon-trout hybrids is not due to alterations in sperm experience via ovarian fluid, and therefore is likely caused by the egg itself.

Due to the diversity of cryptic female choice mechanisms and hybridization across taxa, we recommend conducting other studies in other systems with robust cryptic female choice to fully understand the role females can play in influencing paternal effects. For example, in *Drosophila*, differing placement of sperm in the female’s reproductive tract results in a paternal effect derived from how far the sperm must swim to fertilize the egg (Manier et al., 2013). In black-legged kittiwakes (*Rissa tridactylyus*), females eject older ejaculates in their reproductive tract in favor of newer ejaculates (Wagner et al., 2004), which is known to strongly affect offspring phenotype (White et al., 2008). In these examples sperm experience is likely to drastically vary between males and within ejaculates, which could increase the strength or prevalence of gamete mediated paternal effects.

## Acknowledgments

We thank the staff of the Environmental Resources Management Association for the acquisition of salmon gametes. Assistance in collecting trout gametes and performing the experiment was provided by Terry Sullivan, Madison Philipp, Coady Fitzpatrick, Sydney London, Taylor Hughes, and Alexander Flynn. Funding was provided via Memorial University of Newfoundland, and grants to CFP from the Atlantic Salmon Conservation Foundation, the Natural Sciences and Engineering Research Council of Canada, the Canada Foundation for Innovation, and the Research and Development Corporation of Newfoundland and Labrador. Work was conducted under approved permits from Fisheries & Oceans Canada, and Memorial University of Newfoundland’s Animal Care Committee. We thank Travis van Leeuwen, Peter Westley, Ian Jones and Simone Immler for comments on an earlier version of the manuscript.

---

**Supplement Table 1:** Incubation tubes set up for each treatment in each block until 575 accumulated thermal units (ATUs). Embryos were equalized for density at a maximum of 25 individuals per tube at 575 ATUs. For treatments within blocks with poor hatch success, all embryos were preserved at hatch for timing and size measurements.

| **Tube Number** | **Procedure** |
| --- | --- |
| 1 | Held hatchers until 575 ATUs, no data collected until 800 ATUs |
| 2 | Moved into empty tube 4 at hatch, data collected on hatch timing and size at 800 ATUs |
| 3 | Moved into empty tube 4 at hatch, data collected on hatch timing and size at 800 ATUs |
| 4 | Embryos were split into tubes 1 and 7 at 350 ATUs. Held hatchers from tubes 2 and 3 until 575 ATUs |
| 5 | Preserved at hatch for timing and size data. |
| 6 | Preserved at hatch for timing and size data. |
| 7 | Held hatchers until 575 ATUs, no data collected until 800 ATUs |

## Notes

### Competing Interest Statement

The authors have declared no competing interest.

